# Identification of novel genes in cattle (*Bos taurus*) and biological insights into their function in embryo development

**DOI:** 10.1101/2024.03.15.585311

**Authors:** Gustavo P. Schettini, Michael Morozyuk, Fernando H. Biase

**Author notes:** Corresponding author: West Campus Dr, Blacksburg VA, 24061.

## Abstract

Appropriate regulation of genes expressed in oocytes and embryos is essential for acquisition of developmental competence in mammals. Here, we hypothesized that several genes expressed in oocytes and pre-implantation embryos remain unknown. Our goal was to reconstruct the transcriptome of oocytes (germinal vesicle and metaphase II) and pre-implantation cattle embryos (blastocysts) using short-read and long-read sequences to identify putative new genes. We identified 274,342 transcript sequences, and 3,033 of those transcripts do not match a gene present in an annotation, thus are potential new genes. Notably, 63.67% (1,931/3,033) of potential novel genes exhibited coding potential. Also noteworthy, 97.92% of the putative novel genes overlapped annotation with transposable elements. Comparative analysis of transcript abundance identified that 1,840 novel genes (recently added to the annotation) or potential new genes were differentially expressed between developmental stages (FDR<0.01). We also determined that 522 novel or potential new genes (448 and 34 respectively) were upregulated at eight-cell embryos compared to oocytes (FDR<0.01). In eight-cell embryos, 102 novel or putative new genes were co-expressed (|r|>0.85, P<1×10^-8^) with several genes annotated with gene ontology processes related to pluripotency maintenance and embryo development. CRISPR-Cas9 genome editing confirmed that the disruption of one of the novel genes highly expressed in eight-cell embryos reduced blastocyst development (ENSBTAG00000068261, P=1.55×10^-7^). In conclusion, our results revealed several putative new genes that need careful annotation. Many of the putative new genes have dynamic regulation during pre-implantation development and are important components of gene regulatory networks involved in pluripotency and blastocyst formation.

## INTRODUCTION

Oocytes (the female egg) and pre-implantation embryos express thousands of genes (Driver et al. 2012; Lavagi et al. 2018; Chitwood et al. 2013; Reyes et al. 2015; Robert et al. 2011; Walker and Biase 2020; Graf et al. 2014), and the coordinated regulation of their transcriptional activation or repression is critical for a successful embryo development. Understanding the genetic mechanisms underlying oocyte and embryo development is crucial for unveiling the causes of imbalance that lead to developmental arrest. The assessment of transcriptome profiles using high throughput sequencing has been fundamental in shedding light on gene expression differences between oocyte and embryo stages in cattle (Wrenzycki 2018; Martínez-Moro et al. 2022; Graf et al. 2014). Data from high throughput sequencing have also been used to enhance functional annotation by discovering novel mRNA isoforms and other potential classes of RNA molecules in cattle oocytes and embryos at different developmental stages (Gilchrist et al. 2016; Mondou et al. 2012; Ranjitkar et al. 2023; Wang et al. 2020).

While high throughput sequencing of short-reads provide accurate sequences at massive abundance, their ability to detect long transcripts is limited due to the maximum read length of 150 nucleotides (Stark et al. 2019). In contrast, long-read sequencing technologies such as Pacific Biosciences and Oxford Nanopore Technology (ONT) have demonstrated the capability to generate sequences greater than 10kb and detect full-length transcripts (Wang et al. 2021; van Dijk et al. 2018). By exclusively utilizing ONT long-reads, Halstead et al. (2021) identified several unknown transcripts in 32 tissues from Hereford cattle breed, including tissue-specific isoforms. This discovery unveiled potential unannotated mRNA isoforms and non-coding RNA classes that are missing from the current official cattle annotation and may play key roles in biological processes.

Despite the ability of long-read sequencing technologies such as ONT to detect full-length transcripts, efforts to improve flow cells, chemistry, and basecalling algorithms have increased the accuracy, since these technologies have been prone to higher error rates compared to short-read sequencing methods (Sanderson et al. 2023; Stark et al. 2019). On the other hand, the combination of short-read and long-read sequencing technologies has shown significant advantages in the genome assembly in cattle (Rosen et al. 2020; Chang et al. 2021; Heaton et al. 2021; Ross et al. 2022) and other livestock species (Bickhart et al. 2017; Kalbfleisch et al. 2018; Warren et al. 2017; Warr et al. 2020) since this hybrid approach has facilitated the discovery of new transcripts, identification of structural variants, gap closure, and improved sequence accuracy. Also, it has become the preferred method for reconstructing genomes and transcriptomes in various organisms, as it enables the generation of more accurate and longer contig and scaffold sequences by leveraging both long-read and short-read sequencing data (Sanderson et al. 2023; Wang et al. 2019).

Despite the benefits of combining long and short sequences for transcriptome reconstruction, limited studies have assessed the transcriptome profile of cattle samples using this hybrid approach. Here, we hypothesized that there are several genes expressed in oocytes and pre-implantation embryos that are not yet annotated and many of those genes are functionally important for embryo development. Therefore, we aimed to (a) de novo reconstruct the transcriptome of oocytes and pre-implantation embryos using short- and long-sequences, and (b) evaluate the potential role of those new genes in pre-implantation embryos. A comprehensive analysis of novel genes and putative new genes (not yet annotated) provides new insights into the complex gene regulatory networks at the time of embryo genomic activation and blastocyst formation.

## RESULTS

### Overview of the data sequencing data and hybrid de novo transcriptome assembly

We generated approximately 1.8 billion, 722.49 million, 1.9 billion and 1.05 billion pairs of raw short-reads from 45 oocytes at the germinal vesicle stage (GV), 17 oocytes at the metaphase II stage (MII), 28 embryos at the eight-cell (8c) stage, and 22 embryos at the blastocyst (BL) stage, respectively (Supplemental Table S1). We also produced 23.76 million base-called long-reads (6.03 – GV, 6.01 – MII, and 11.72 – BL) at an average length of 1,417 nucleotides long (minimum: 20; maximum: 91,586).

A flow chart with the schematics of the study is depicted in Figure 1. For transcriptome assembly, we grouped all short-read sequences for alignment, quality filtering, and coverage normalization (10-30x range). These processes resulted in 41.66 million short pair-end reads used as input along 23.76 million long sequences into RNAspades assembler (Prjibelski et al. 2020). The resulting transcriptome assembly generated 277,818 sequences with a mean length of 2,462 nucleotides (min.: 77; max.: 93,634) (Supplemental Table S2). After aligning to the genome with GMAP (Wu and Watanabe 2005) we obtained the coordinates for 274,342 transcripts (Table 1).

**Figure 1.**
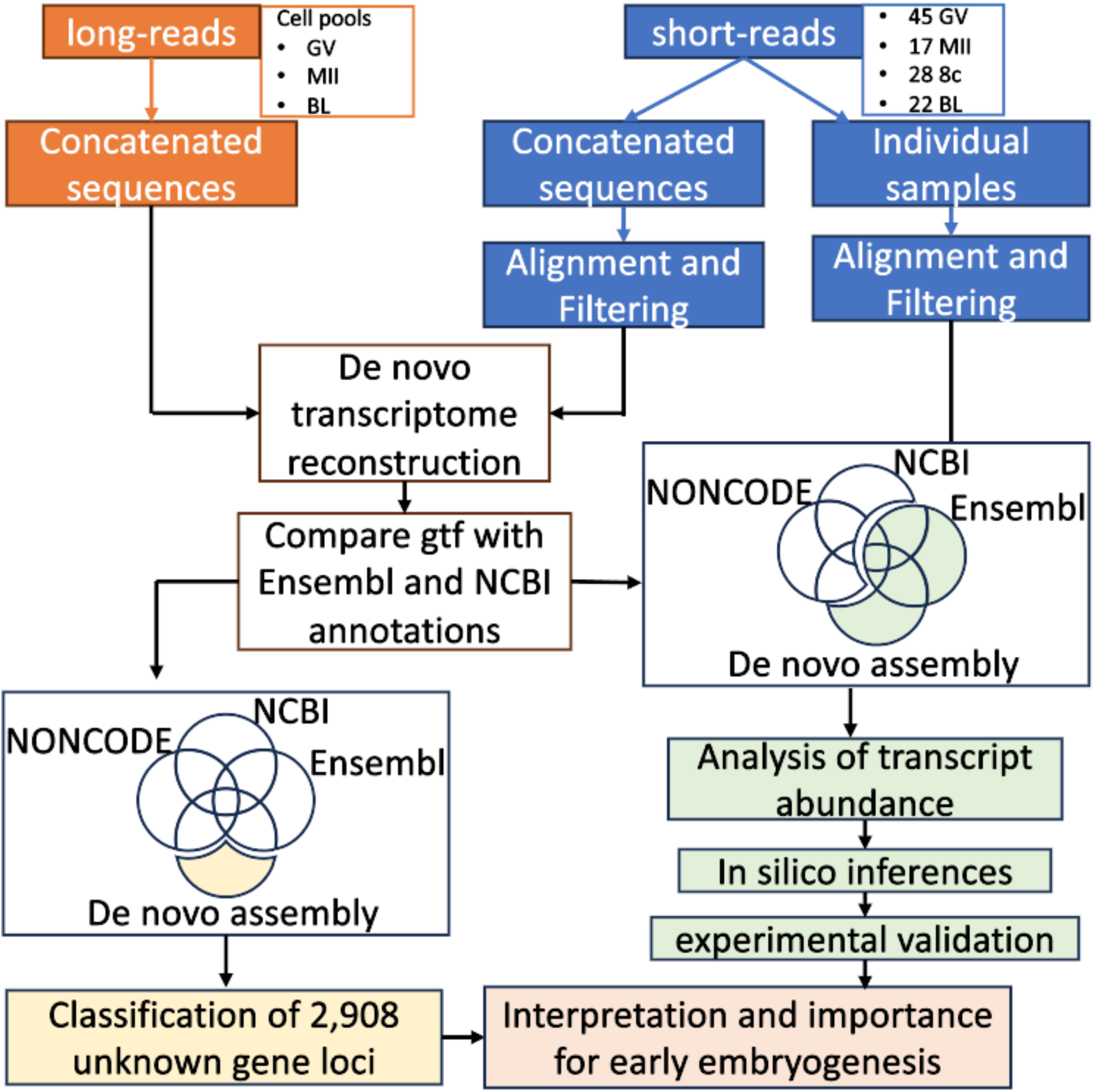
Overview of bioinformatic steps for transcriptome reconstruction and identification of unknown gene loci, followed by categorization and functional categorization.

**Table 1.** Summary of de novo transcriptome reconstruction and mapping to annotation.

|  | Total number of genomic regions | Novel genes |
| --- | --- | --- |
| De novo assembly | 274,342 | - |
| Ensembl | 202,945 | 7,990 |
| NCBI/RefSeq | 10,117 | 1,062 |
| lncRNA (NONCODEv5) | 882 | - |
| Not yet annotated-current study | 22,305 | 3,033 |

### Novel gene loci identified in oocyte and pre-implantation embryos

After comparisons with reference annotation information from Ensembl (ARS-UCD1.3.111), NCBI (ARS-UCD.2.0.GCF_002263795.3-RS_2023_09), and lncRNA NONCODEv5 databases (Figure 1), we identified 22,305 transcripts that did not overlap coordinates with the annotations assessed. Those transcripts were reduced to 3,033 loci (see methods for details) that have not yet been annotated, which we referred to as potential novel genes (Table 1).

Based on homology with sequences present in the NCBI database, we determined that 63.67% (1,931/3,033) of the putative novel genes have coding potential, while there was an indication that the remaining 36.33% putative novel genes are transcribed as long non-coding RNAs (Table 2). The majority (97.92%, 1,926 coding; 1,044 noncoding) of the putative novel genes had overlapping coordinates with at least one transposable element (Table 2). There is a greater (99.74%) proportion of potential new genes with coding potential associated with transposable elements relative to the annotated novel genes (93.12%, *X*^2^=123.99, P < 2.2^-16^, 2-sample test for equality of proportions). This distinct pattern of colocalization with a transposable element is less prominent for non-coding potential new genes (94.74%) relative to annotated new genes (93.12%, *X*^2^=3.68, P=0.03, 2-sample test for equality of proportions).

**Table 2.**
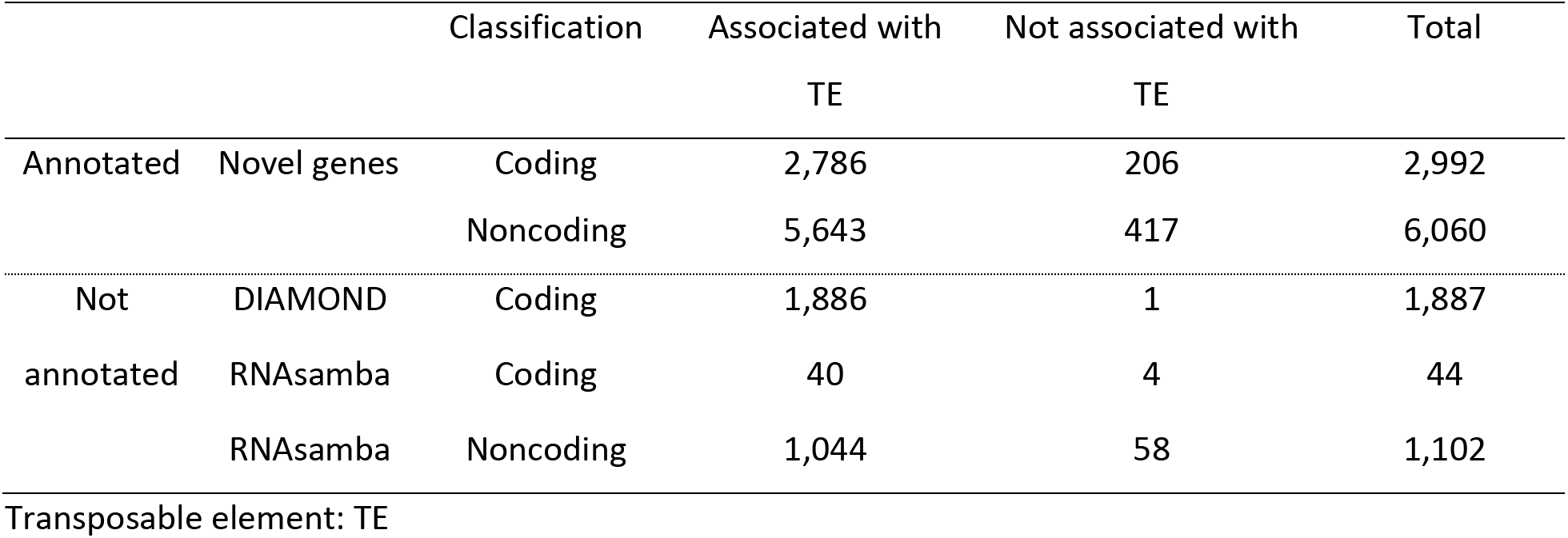
Summary of classification of annotated novel genes and potential novel genes.

### Expression of novel and putative novel loci during pre-implantation development

After filtering lowly expressed genes (<5 counts per million, CPM and < 15 samples), we retained 12,932 annotated protein-coding or lncRNA genes, 1705 annotated novel protein-coding or lncRNA genes and 294 potential new genes (out of 3,033 shown on Table 1), for which we estimated robust relative transcript abundance. We noted that the potential new genes were distributed across all chromosomes (Figure 1 A). All three gene subsets produced similar global separation of samples, including a similar pattern of dispersion of samples, with the broadest dispersion observed among embryos collected at the 8-cell stage (Figure 2 B, C, D). From a global perspective, novel annotated and putative new genes had similar patterns of expression, with an overall less transcript abundance across oocytes or embryos, relative to the annotated genes (Figure 2 E, F, G).

**Figure 2.**
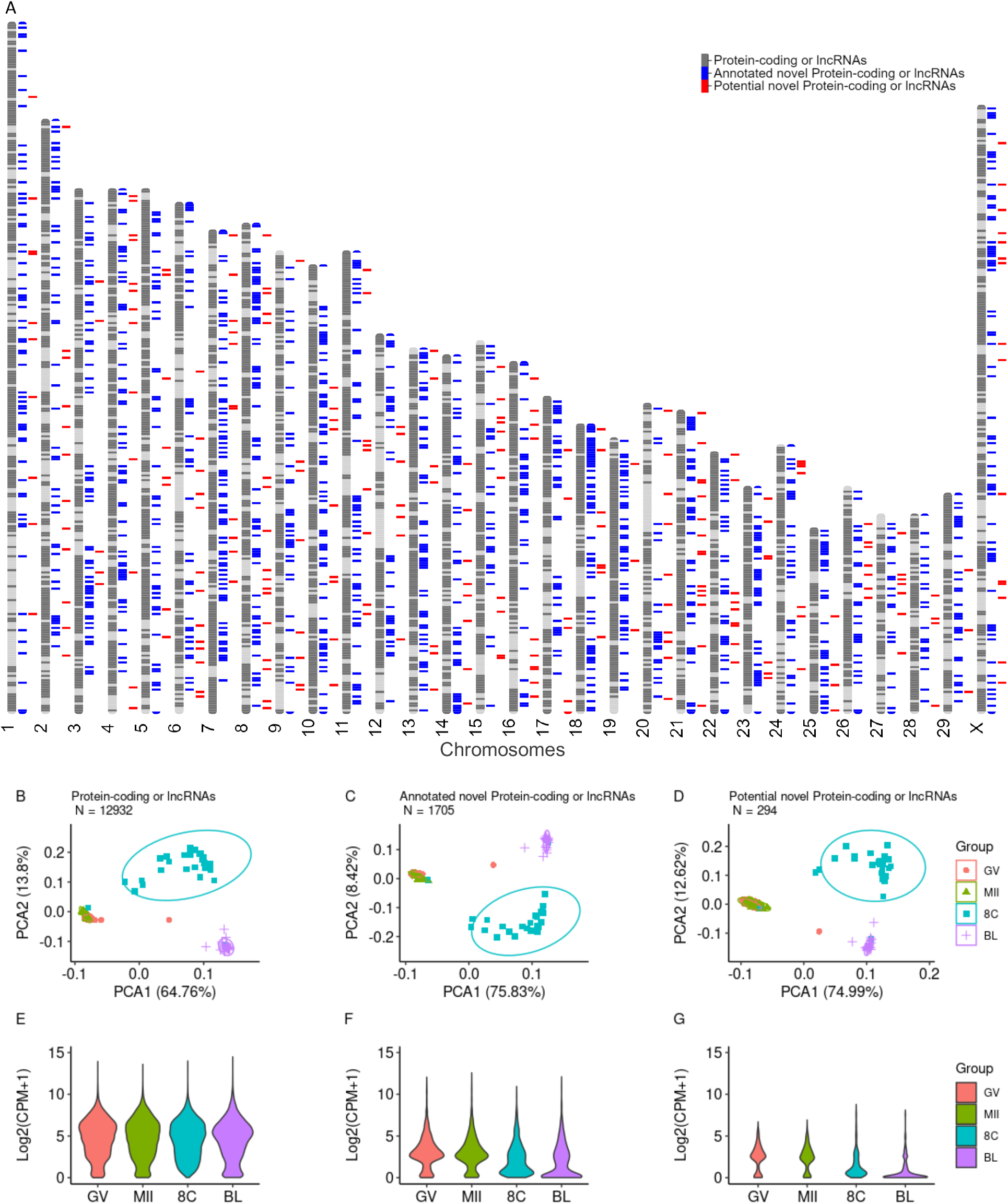
Overview of the transcriptome (protein-coding or long-noncoding genes) of single oocytes or pre-implantation embryos. (A) Distribution of the novel (blue) and potential new genes (red) across the cattle genome. Principal component analysis of annotated (B), annotated as novel (C) and putative novel genes (D). Distribution of the average transcript abundance of annotated (E), annotated as novel (F) and putative novel genes (G).

In order to understand the pattern of regulation of the novel and potential novel genes during pre-implantation development, we focused our analysis of differential transcript abundance on the 1999 loci that are novel and annotated (1705) or potential novel genes (294) (Figure 3, Supplemental Table S3-S8). Most (92%) of the loci showed alteration in transcript abundance between two developmental stages. Not surprisingly, comparisons between oocytes and embryos (eight-cell or blastocyst) resulted in a greater number of loci with differential transcript abundance (Figure 3A). On the other hand, it was interesting to observe that 312 loci had differential transcript abundance between eight-cell and blastocyst stages (Figure 3A).

**Figure 3.**
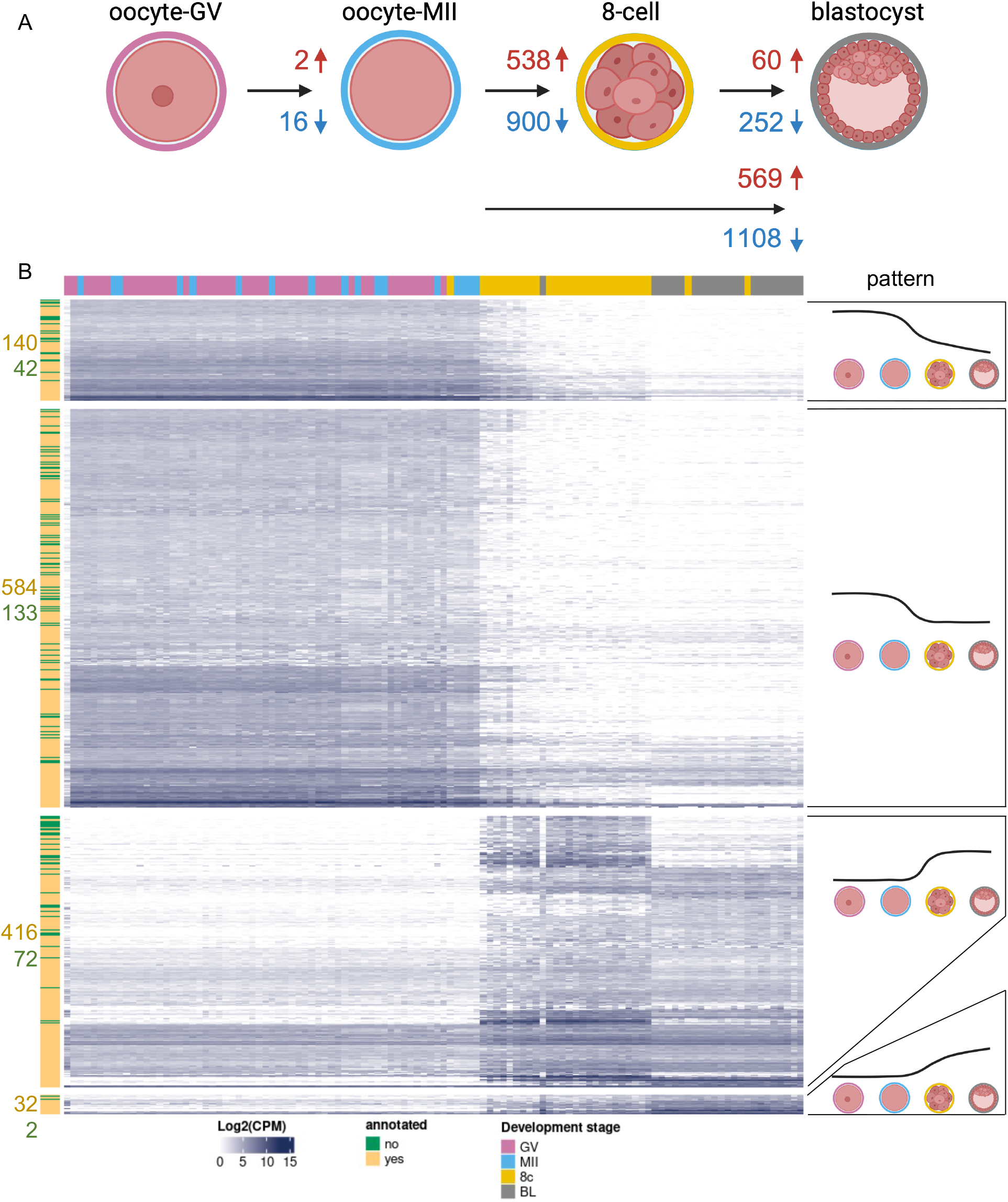
Differential transcript abundance of 1999 novel or putative novel loci during pre-implantation stages. (A) Summary of the number of loci with differential transcript abundance between two stages (|Log2(FC)|>1 and FDR <0.01). (B) Patterns of differential transcript abundance between oocytes and embryos. Only loci with significant differential transcript abundance based on contrasts between contiguous stages are represented (i.e.: MII oocyte versus GV oocyte, eight-cell embryos versus MII oocyte, blastocysts versus eight-cell embryos)

Changes in transcript abundance during the pre-implantation development revealed that several genes followed four distinct patterns of expression (Figure 3B). The most common pattern (717 loci) was the depletion (FDR<0.01) of transcripts between oocytes and eight-cell stage followed by a non-significant change in transcript abundance between eight-cell and blastocyst stage. Next, 488 loci showed an increase (FDR<0.01) in transcript abundance between oocytes and eight-cell stage followed by a non-significant change in transcript abundance between eight-cell and blastocyst stage. The other two patterns were the continuous depletion or increase of transcript abundance between oocytes, eight-cell and blastocyst stage (182 and 34 loci respectively, FDR<0.01).

### Functional characterization of novel and putative novel loci in pre-implantation embryos

To better understand the function of novel annotated genes and potential new loci in pre-implantation embryo development, we conducted a co-expression analysis between 552 loci that showed an increased transcript abundance in embryos relative to MII oocytes (3^rd^ and 4^th^ patterns in Figure 3A) and the 12,932 protein-coding and long non-coding genes that have been annotated. We also focused the analysis on eight-cell and blastocyst embryos.

In eight-cell embryos, there were 247 novel annotated genes and 41 putative new loci showing significant co-expression with 2761 annotated genes (|r|>0.85, P≤ 1×10^-8^, Figure 4A, Supplemental Table S9). Interestingly, biological processes “translation” (167 genes), “ribosomal small subunit biogenesis” (33 genes) and “rRNA processing” (42 genes) were enriched among the 2761 co-expressed genes (FDR<0.01, Supplemental Table S11). In blastocysts, there were 157 novel annotated genes and 23 putative new loci showing significant co-expression with 1667 annotated genes (|r|>0.85, P≤ 5×10^-7^, Figure 4B). Those 1667 showed enrichment for “translation” (154 genes), “ribosomal large subunit biogenesis” (24 genes), “ribosome biogenesis” (29 genes), “rRNA processing” (33 genes), “ribosomal small subunit biogenesis” (27 genes) and “RNA splicing” (36 genes) biological processes (FDR<0.01, Supplemental Table S12).

**Figure 4.**
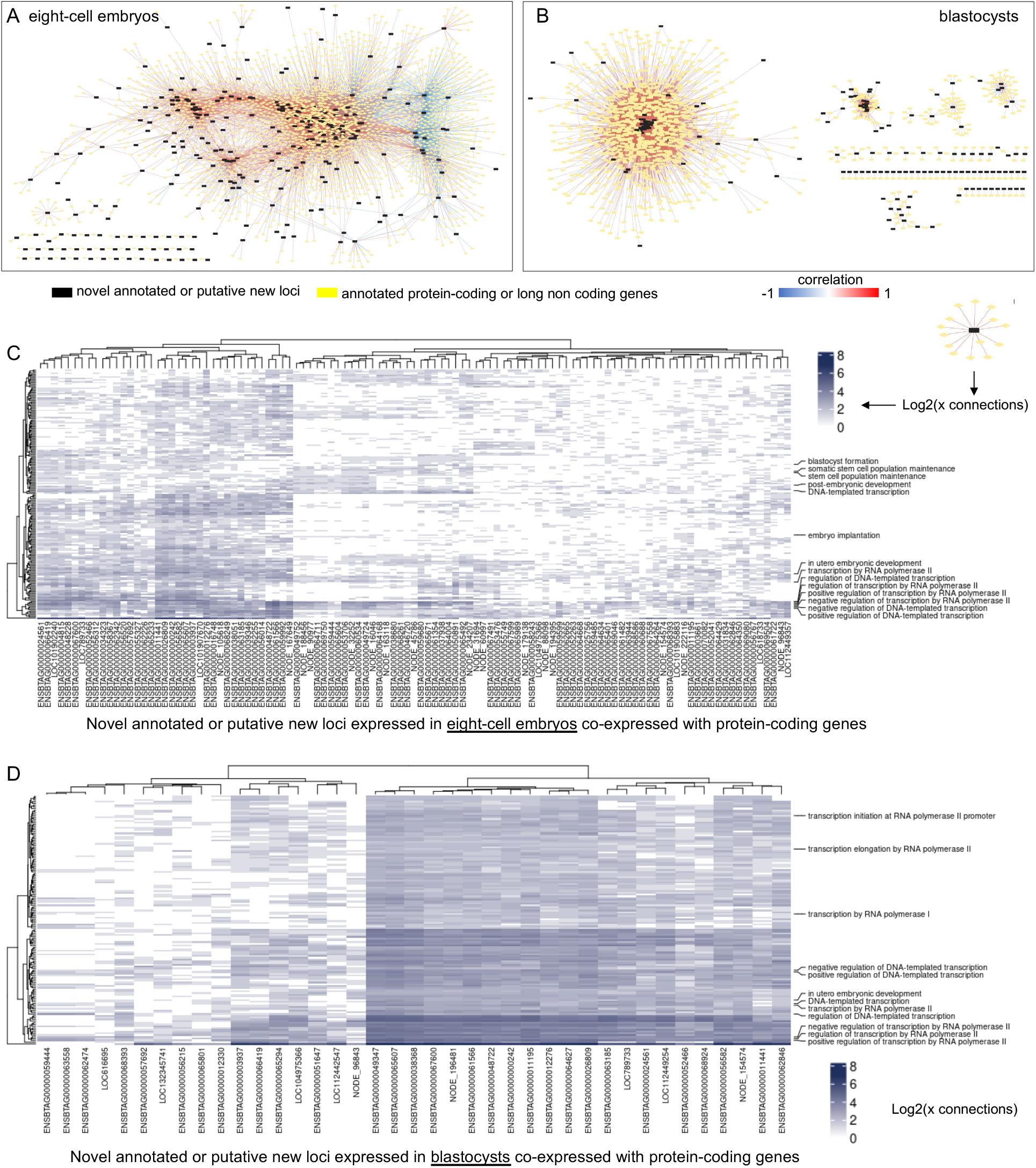
In silico functional characterization of the novel annotated genes and putative new loci. Co-expression networks with annotated protein-coding genes and long non-coding genes for eight-cell stage (A) and blastocysts (B). Heatmaps of connectivity for eight-cell embryos (C) and blastocysts (D) based on the co-expression networks with genes that are functionally annotated in gene ontology database.

A careful interrogation of the genes co-expressed with novel or putative new loci also revealed that many novel genes were highly correlated with several genes involved in the regulation of transcription in both eight-cell and blastocyst stages (Figure 4 C, D). Most notably, at eight-cell stage, 22 putative new genes and 80 novel annotated genes showed co-expression with genes (*BRAF*, *BYSL*, *EIF4ENIF1*, *ERRFI1*, *FOXO3*, *FUT10*, *GABPA*, *GNL3*, *IGF2BP1*, *KLF4*, *KLF10*, *LEO1*, *MYC*, *NANOG*, *NFIB*, *PAF1*, *PCM1SUPT6H*, *PRDM14*, *PROX1*, *RIF1*, *RBPJ*, *RPL7L1*, *RRP7*, *SIRT6*, *SF3B6*, *SOX4I, STAT3*, *TBX3*, *TEAD4*, *TPT1*, *TRIM28*, *VMP1*, *WDR74*, *WDR43*, *ZP3*, and *ZHX2*) that are annotated with biological processes extremely relevant for embryo development (“blastocyst formation”, “stem cell population maintenance” and “embryo implantation”, Figure 4 C). These co-expressing pairs of genes were not present in blastocysts (Figure 4 D).

We selected the novel gene ENSBTAG00000068261 for further validation via CRISPR-Cas9D10A approach. This gene was selected because of its abundant transcription in eight-cell embryos (Figure 5A) and co-expression with genes associated with “cell differentiation” (*ALKBH1*), “blastocyst formation” (*SUPT6H*), “regulation of blastocyst development” (*KLF4*), “regulation of transcription by RNA polymerase II” (*KLF4*, *KLF17*, *SNAI1*, *ZNF394*, *ZNF570*, *ZNF608*, *ZIM3*, *ZSCAN4*), and “stem cell population maintenance” (*KLF4*). The introduction of ribonucleoproteins targeting exon 2 of ENSBTAG00000068261 into zygotes caused deletions in the targeted sequence (Figure 5 B). Embryos subjected to editing displayed similar cleavage rates when compared to controls (ribonucleoprotein with scrambled guide RNA), accessed at ∼45 hours post fertilization (hpf, 70.5%±12.7 versus 74.2%±6.09; P = 0.531). By contrast, there was a significant reduction in blastocysts developed relative to controls at ∼168hpf (7.52%±4.57 versus 19.8%±4.57; P = 1.57×10^-3^) and ∼190hpf (11.4%±6.32 vs. 36.3%±6.85; P = 1.55×10^-7^)(Supplemental Tables S13-14). A greater proportion of embryos arrested development at the morula stage (Figure 5 C).

**Figure 5.**
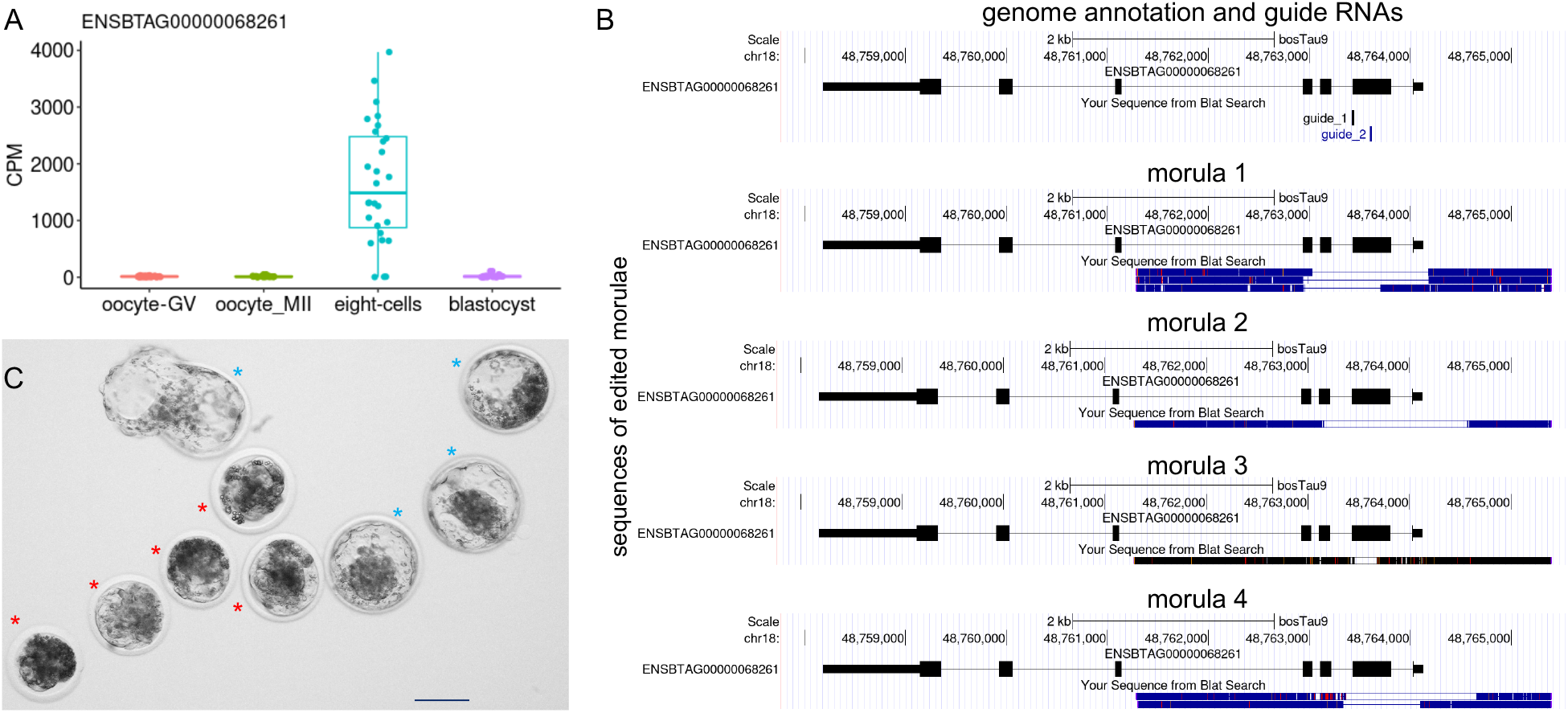
Gene knockout to evaluate the importance of novel genes in pre-implantation development. (A) Transcript abundance of the novel gene ENSBTAG00000068261 in individual oocytes, eight-cell embryos and blastocysts. (B) Examples of sequences containing deletions of exon 2 of the ENSBTAG00000068261 novel gene. (C) Representative images of the embryos that arrested development at the morula stage (red asterisk) compared to control blastocysts (blue asterisk, all collected and imaged ∼190hpf). Scale bar represents 100µm.

## DISCUSSION

Our driving motivation was to determine whether there are genes that have yet to be identified in the cattle genome. Here we carried out a hybrid transcriptome reconstruction using high-throughput sequencing data collected from oocytes and early developing cattle embryos. We report 3,033 potential new genes with no annotation in major genomic databases (Ensembl (Flicek et al. 2014), NCBI/RefSeq (Sayers et al. 2022) or NONCODEv5 (Fang et al. 2018)) along with 9,052 loci recently (January/2024) added to Ensembl and RefSeq annotations. It was notable that a set of newly identified genes can integrate gene regulatory networks with genes involved in biological functions essential for embryo development. We also demonstrated the importance of those new genes by disturbing its expression in pre-implantation embryos, which led to developmental arrest at the morula stage. The results confirm our hypothesis and reveal an important gap in our understanding of the genome function in the early stages of development.

The identification of 3,033 potential new genes may seem high, however, congruent lines of evidence indicate that the results are not artifacts. First, the use of a hybrid data, combining short and long-reads, is beneficial for identifying new transcripts (Halstead et al. 2021) and produces contigs in *de novo* assemblies with high confidence (De Maio et al. 2019; Goodwin et al. 2015). The number of contigs that we report were also filtered by those that contained a minimum of short reads mapping to them, thus increasing our confidence of detection. Second, in the process of concluding our study, a new annotation was released in January/2024 and 2238 transcripts or contigs that were initially categorized as potential new genes were reassigned to novel genes (see Table 2). Thus, the most recent annotation from Ensembl or NCBI corroborated 42% of our initial transcripts (2238/5271). Third, beyond bioinformatic predictions conducted by NCBI and Ensembl, the samples we used in our study (oocytes and blastocysts) are not commonly included in similar research previously executed (Goszczynski et al. 2021; Veselovska et al. 2015; Halstead et al. 2021). Even in our samples, the putative new genes were not expressed at high levels. Fourth, the pattern of expression of those potential new genes, discussed further below, is indicative of functional units in the genome (Gerstein et al. 2007; Goldman and Landweber 2016). Thus, the findings indicate that the new transcripts identified in our de novo transcriptome assembly are from putative new genes.

Since there was limited information related to the recently identified new genes and no data related to potential new genes, we performed a classification step and identified that most of the gene loci were associated with at least one TE by coordinate overlapping. Such a high number of associations could be explained, first, by the high TE activity and expression during germline and pluripotent-like cells, such as oocytes and embryos in mammals (de la Rosa et al. 2024; Gifford et al. 2013), which are suggested to play important roles in pluripotency maintenance (de la Rosa et al. 2024) and transcriptional regulation (Fueyo et al. 2022; Gebrie 2023). Second, TE are not randomly distributed in the genome, and it has been co-located with more active genes along with developmental stages (Zhang and Mager 2012; Gebrie 2023). Notwithstanding, TE have been involved in the origin of new lncRNA (Johnson and Guigó 2014; Kelley and Rinn 2012), and have been identified in several protein-coding sequences (van de Lagemaat et al. 2003; Nekrutenko and Li 2001; Brandt et al. 2005), which supports our findings.

Several of the novel genes and possible new genes had significant differential transcript abundance in eight-cell embryos relative to oocytes. This finding is similar to patterns previously identified in the literature considering that a major embryo genome activation occurs in cattle at the 8-cell stage (Ming et al. 2020; Graf et al. 2014; Jiang et al. 2014). The transcriptional activation of novel genes and possible new genes at the eight-cell stage supports the fact that those loci are actual genes and the rules of their regulation follow the annotated protein-coding and long non-coding genes.

The increase in transcripts at the eight-cell stage is a strong indication of their importance for major events that follow the embryo genome activation. To that end, the fact that most of the annotated co-expressing genes were related to translation, indicates that hundreds of those uncharacterized genes produce proteins that either have a direct role in protein synthesis or participate in the regulation of downstream genes directly related to protein synthesis. The notion that those uncharacterized genes participate in the regulation of other genes is also supported by the co-expression of hundreds of uncharacterized genes with genes annotated with transcription related to biological functions. Another in silico layer of support for the importance of those novel genes and possible new genes (522 genes) was their co-expression with several genes annotated with biological functions related to embryo development and/or stem cell pluripotency, such as *KLF4* (Halstead et al. 2020; Wei et al. 2009, 2013)*, ALKBH1* (Pan et al. 2008; Ougland et al. 2016), and *SUPT6H* (Bedi et al. 2015; Oqani et al. 2019). Collectively, the increase in transcription at the eight-cell stage and co-expression with genes that are important for embryo development support their critical role in the early stages of development.

The different expression patterns of novel genes and potential new genes during early embryonic development stages indicate a stage-specific importance that could also be associated with the TE composition, in which families and sub-families are suggested to be expressed pluripotency-stage-specifically (Li et al. 2023; Todd et al. 2019). Such a relationship is also suggested due to the presence of the same transcription factor binding sites located by TE (Glinsky 2015; Testori et al. 2012), which explains the high correlation with genes enriched in transcriptional and translation regulation, and more specifically, stem cells population maintenance and embryo/blastocyst development, such as *STAT3, PRDM14*, and *NANOG*, which are well-known to be involved in pluripotency and embryo development in cattle (Wei et al. 2017; Meng et al. 2015; Khan et al. 2012; Sang et al. 2020).

In order to test how critical those novel or potential new genes are for embryo survival, we deleted exon 2 of the gene ENSBTAG00000068261, also identified as LOC132342749 on RefSeq/NCBI database. This gene was originally selected for gene editing as a potential new gene but was annotated as a novel gene in January/2024, receiving the description of “F-box only protein 27-like”. As a member of the F-box protein genes, it is possible that F-box only protein 27-like participates in the SFC complex, which is comprised of Cullin, RBX1, and SKP1 proteins, all of which have an important role in the ubiquitination of maternal proteins during the maternal-to-zygotic transition (Xie et al. 2019). The SFC complex also participates in the regulation of cell proliferation and differentiation (Skaar et al. 2013; Randle and Laman 2016), which is aligned with the co-expression between transcripts of F-box only protein 27-like and other genes known to regulate pluripotency. A role of F-box only protein 27-like in pluripotency is also supported by embryonic arrest at the morula stage in knockout embryos. A Similar phenotype was observed by Kinterova et al. (2019) when SFC complex activity was inhibited. The findings are an example that several genes that remain poorly studied have important biological implications in early embryo development.

Fifteen years after the release of the first draft of the cattle genome (Elsik et al. 2009), our study reveal that thousands of genes have yet to be annotated. The findings also provide multiple lines of evidence that possible new genes have a critical role in early stages of embryo development, some of which are necessary for the correct formation of blastocysts. The findings herein add important insights to the complex biology involving genes expressed in oocytes and embryos and their role in the acquisition of developmental competence to achieve a successful pregnancy.

## METHODS

No live animal was handled for this work, thus approval by the Institutional Animal Care and Use Committee was not necessary.

### Sample collection

All procedures for oocyte maturation, embryo culture and media composition were described in detail elsewhere (Ortega et al. 2017; Nix et al. 2023a; Tríbulo et al. 2019). *Bos taurus* ovaries were collected from a commercial slaughterhouse and transported to the laboratory in a 0.9% saline solution and antibiotic and antimycotic solution (1x, ThermoFisher Scientific, Waltham, MA, USA). First, ovarian follicles were slashed, and the cumulus-oocyte complexes (COCs) were retrieved from the follicular fluid into oocyte collection media (OCM). Next, we assessed the morphological properties of the COCs under a stereoscope and only COCs presenting homogeneous ooplasm and more than three layers of compact cumulus cells were selected for sample collection or *in vitro* maturation (IVM) at the density of ten COCs in 50μl of IVM medium in a humidified incubator at 5% CO^2^ and 38.5°C for 22-24 hours.

For in vitro fertilization (IVF), we washed in vitro matured COCs trice in 4-(2-hydroxyethyl)-1-piperazineethanesulfonic acid (HEPES) buffered synthetic oviductal fluid (HEPES-SOF), followed by two washes in fertilization medium (SOF-Fert). The IVF was set up in SOF-Fert with the addition of sperm (1,000,000 sperm/ml) and incubation for 16-18 hours in a humidified incubator at 5% CO_2_ and 38.5°C. In vitro culture (IVC) of putative zygotes was conducted by removal of cumulus cells adhered to the zona pellucida in HEPES-SOF, which was followed by three washes in SOF culture media (SOF-BE). Then, we placed 25-30 presumed zygotes, in micro drops (50µl) of culture media covered by mineral oil and in a humidified incubator at 38.5 °C under 5% CO_2_, 5% O_2_, and 90% N_2_.

We collected oocytes at the germinal vesicle stage and metaphase II stage by washing the COCs in Trypsin (TrypLE^TM^ Express, gibco, Grand Island, NY) twice followed by a 10-minute incubation. We removed the remaining cumulus cells adhered to the zona pelucida by repeated pipetting. Then, we washed the oocytes in phosphate buffered saline solution (PBS, ThermoFisher Scientific, Fair Lawn, NJ). Embryos were collected at either 60 hours post culture (at 8-cell stage) or 164 hours post culture (at blastocyst stage) and washed twice in PBS. Oocytes were collected either individually in 1µL of PBS or in groups of 50 in 5 µL of PBS. Embryos were collected either individually in 1µL of PBS or in groups of 25 embryos in 5 µL of PBS. All samples were preserved at -80°C until RNA extraction.

### RNA extraction and Hight-throughput sequencing

We extracted total RNA from all samples using TRIzol reagent (Rio et al. 2010) with Phasemaker Tubes using the procedures described elsewhere (Biase 2021), followed by storage at -80°C until further processing.

To conduct short read high throughput sequencing, we prepared libraries of single oocytes or single embryos (45 oocytes at the GV stage, 17 oocytes at the MII stage, 28 8-cell embryos, and 22 blastocysts) using a procedure described elsewhere (Bagnoli et al. 2018; Biase 2021). Briefly, RNA pellets were resuspended into a solution containing oligo-dTVN oligonucleotide (1mM) (Supplemental Table S15) and heated in a thermocycler at 72°C for three minutes. Reverse transcription was carried out by adding 5µl of a solution containing 200 U/µl Maxima H Minus Reverse Transcriptase, 1x Maxima RT Buffer, 7.5% PEG 8000, 10 mM dNTPs, and 2 µM of a template-switching oligo (Supplemental Table S15) and to the RNA oligonucleotide mix for 1h 30min at 42°C. Next, cDNA was purified by AMPure XP beads.

Promptly after cDNA purification procedures, an amplification mix containing 1.25U Terra polymerase, 1x Terra direct buffer, and 0.1µM of cDNA amplification primer (Supplemental Table S15) was added to each sample tube. We used purified cDNA for amplification by polymerase chain reaction (PCR) under the following conditions: 98°C for 15 minutes, 68°C for 5 minutes, and 72°C for 10 minutes, and presumed at 8°C after the last cycle. The amplification products were cleaned using AMPure XP magnetic beads, quantified with a Qubit 4.0 fluorometer, and quality-assessed by 2100 Bioanalyzer and the Agilent High Sensitivity DNA kit.

According to the manufacturer’s instructions, we used 1ng of cDNA as a template for library preparation with the Nextera DNA Flex Library Prep kit. Followed by PCR amplification (13 cycles) and a purification step using AMPure XP beads. We quantified the libraries using a Qubit 4 fluorometer and assessed their quality using the Agilent High Sensitivity DNA kit in a 2100 Bioanalyzer. Libraries were sequenced at Vanderbilt Technologies for Advanced Genomics at Vanderbilt University – VANTAGE to produce 150 bp pair-end reads in a HiSeq 2500 or NovaSeq 6000 Illumina sequencer (Illumina, Inc. San Diego, CA).

To proceed with high throughput sequencing of long reads, we extracted RNA from pools prepared for each development stage (GV, MII, and BL, N = 50 in each pool). Next, we amplified the material with TeloPrime Full-Length cDNA Amplification Kit V2 (Lexogen, Vienna, Austria), following the manufacturer’s procedures and 20 PCR cycles. Next, the libraries were prepared with the Native Barcoding kit 24 V14 (Oxford Nanopore Technologies, Oxford, England). We sequenced the libraries using the MinION Mk1C sequencer with R9.4.1 (GV oocytes) and R10.4.1 (MII oocytes, and BL) flow cells (ONT Ltd., Oxford, United Kingdom).

### Pre-processing RNA-seq sequences

We trimmed the adaptors from short-reads sequences produced from 45 GV, 17 MII, 28 8c, and 22 BL samples with Trimmomatic v.0.39 (Bolger et al. 2014) and aligned to the cattle genome (ARS-UCD 1.2) (Rosen et al. 2020) downloaded from the Ensembl database (Flicek et al. 2014), using HISAT2 v2.1.0 aligner (Kim et al. 2019). Next, we used SAMtools v1.10 (Li et al. 2009) and biobambam2 v2.0.95 (Tischler and Leonard 2014) to remove unmapped reads, secondary alignments, PCR duplicates, and duplicated sequences. Later, we converted those filtered alignment files (.bam) to fastq files using the *bomtofastq* command built on the biobambam2 v2.0.95 (Tischler and Leonard 2014). Then, aiming to increase specificity, we concatenated and pre-processed all sequences on BBTools (Bushnell, 2018) to maintain regions with coverage between 10x and 30x coverage.

We conducted base calling of ONT long-reads using Guppy v.6.4.2 (Wick et al. 2019) with the super accuracy algorithm (*dna_r10.4.1_e8.2_260bps_sup.cfg*) and used in the *de novo* assembly.

### Hybrid de novo transcriptome assembly

We conducted transcriptome assembly based on short and long sequences using the hybrid *de novo* assembler rnaSPAdes v3.14.1 (Prjibelski et al. 2020). The *de novo* assembly by RNAspades relies mainly on the SR sequences and only uses the LR sequences as support to close gaps between SR contigs (Prjibelski et al. 2020).

### Unknown or novel gene loci identification

First, we aligned the sequences obtained from the *de novo* assembly to the *Bos taurus* genome (ARS-UCD 1.2) using the GMAP aligner (version 2021-12-17) (Wu and Watanabe 2005) to produce a preliminary annotation (.gff3) and alignment (.sam) using the following parameters to improve accuracy and report only the best sequence paths (*--microexon-spliceprob=1 --nofails --quality-protocol=illumina - -suboptimal-score=0.99 --min-identity=0.90 --npaths=1*). Sequence alignments with more than ten mismatches were filtered out.

Second, we compared our assembly to the Ensembl annotation file (ARS-UCD 1.3.111) with gffcompare v0.12.6 (Pertea and Pertea 2020) to identify transcripts not yet present in the Ensembl annotation. Only transcripts classified with the flag “-u” (unknown) were retained in our annotation file.

The fasta sequences from transcripts listed as unknown relative to the Ensembl annotation were aligned to the NCBI (Sayers et al. 2022) genome (ARS-UCD 1.2) and mapped to the RefSeq annotation (GCF_002263795.3-RS_2023_09). Only transcripts classified with the flag “-u” (unknown) were retained in our annotation file.

Third, we compared the transcripts identified as not present in both Ensembl and NCBI databases to the lncRNA database (NONCODEV5) (Fang et al. 2018). Lastly, we filtered out any unknown transcript located within five kilobases to the boundaries of known genes, containing only one exon and those with less than 300nt length from our annotation file by gffread v0.12.8 (Pertea and Pertea 2020) and GenomicRanges R-package (Lawrence et al. 2013).

We used the annotation file (unknown transcripts) to count short-reads by FeatureCounts (Liao et al. 2014). We retained transcripts with abundance greater than two counts per million (CPM) in five or more samples. Subsequently, we used gffread (Pertea and Pertea 2020) and bedtools (Quinlan and Hall 2010) to reduce the redundancy of our assembly by merging transcripts into unknown gene loci allowing a maximum gap of 200 nucleotides.

### Classification of Unknown loci

In order to classify the unknown gene loci identified in our study, we assessed the coding potential based on high similarity with non-redundant protein (nr) sequences deposited in NCBI database v5 (ftp.ncbi.nlm.nih.gov/blast/db/v5/FASTA - updated on 07/Feb/2024) via DIAMOND v.2.0.11.149 (Buchfink et al. 2021). We used database for potential protein family members located in different regions in cattle (*Bos taurus; Bos indicus; Bos taurus x Bos indicus*), followed by orthologous proteins in the mammalian class, prioritizing human (*Homo sapiens*) and mouse (*Mus musculus*). First, we carried out a local alignment via DIAMOND using parameters to increase specificity, such as an E-score threshold of 1×10^-6^, >90% of subject coverage, and a percentage of identity greater than 90% to report a potential hit for cattle. Second, the remaining loci with no match went through a local alignment to identify potential orthologs in mammalian organisms. The parameters were similar, except for the percentage of identity threshold, which was set at >70%. Unknown gene loci that remained without classification were assessed by a neural network classification model, RNAsamba (Camargo et al. 2020), and classified for coding potential on open read frame (ORFs) and untranslated region (UTRs) features. Lastly, we mapped the unknown loci to known transposable elements (TE) by coordinates overlap conducted by GenomicRanges (Lawrence et al. 2013). TE coordinates were obtained from RepeatMasker (Chen 2004) and retrieved from the University of California Santa Cruz (UCSC) database (Kent et al. 2002; Karolchik et al. 2004).

### Unknown/novel gene loci abundance and differential expression

We determined loci transcript abundance by counting short-reads mapped to the gtf files obtained from Ensembl (ARS-UCD 1.3.111) and NCBI (GCF_002263795.3-RS_2023_09) and our gtf file with unknown loci via FeatureCounts (Liao et al. 2014). Subsequently, we combined the gene raw count matrices and removed NCBI genes with mapped identifiers in the Ensembl annotation.

We normalized libraries using the trimmed mean of M values (TMM) (Robinson and Oshlack 2010), followed by a count per million (CPM) using the edgeR package (Robinson et al. 2010). Loci with less than five CPM in at least 15 samples were filtered out. Then, we conducted the differential gene expression analyses using EdgeR (Robinson et al. 2010) and DESeq2 (Love et al. 2014) R-packages, and loci classified as statistically differentially expressed (DE) in both algorithms if |LogFC| > 1 and FDR < 0.01.

### In silico functional characterization novel or unknown loci

Focusing on pre-implantation embryos, we performed a co-expression analysis on novel genes/potential novel gene loci used for differential expression analysis. The same normalized matrices obtained previously were transformed using an inverse hyperbolic sine (Johnson 1949), followed by Pearson correlation coefficient calculation between the novel genes/potential novel gene loci and Ensembl known genes using WGCNA R-package function *corAndPvalue* (Langfelder and Horvath 2008). We retained absolute correlation values ≥ 0.85 with a P ≤ 0.00005. Further, we conducted a functional enrichment analysis using GOseq R-package (Young et al. 2010) on the genes highly co-expressed, and only biological processes (BP) with more than four genes and FDR ≤0.01 using the Holm–Bonferroni method considered significantly enriched. Additionally, we functionally characterize the novel genes/potential novel gene loci based on co-expressed annotated gene functional information retrieved from biomaRt (Durinck et al. 2009). Biological processes containing > 55 co-expressed genes and in at least 50 novel genes/potential novel gene loci correlations were maintained. Finally, co-expression networks were generated by Cytoscape (v.3.10.1) (Shannon et al. 2003).

### Functional characterization using CRISPR-CAS9

We designed gRNAs (Supplemental Table S15) to target exon 2 of the novel gene locus (ENSBTAG00000068261/LOC132342749) located on chr18:48758182-48764129 using the CRISPOR (Concordet and Haeussler 2018). All gRNAs were purchased as a single RNA molecule (sgRNA) comprising crRNA and transacting crRNA (tracerRNA) from IDT (Integrated DNA Technologies, Research Triangle Park, NC, USA), as well as CRISPR-Cas9D10A nickase V3.

We mixed CRISPR-Cas9D10A and sgRNAs for the formation of ribonucleoproteins in OptiMEM reduced serum medium (Thermo Fisher Scientific, Grand Island, NY) at room temperature for at least 1h before electroporation. The concentrations in the solution for the formation of RNPs were 800ng/µl Cas9D10A and 800ng/µl of each sgRNA. We electroporated the presumptive zygotes (PZ) following the procedures detailed elsewhere (Nix et al. 2023b). We removed the cumulus cells from the PZs and electroporated them in Opti-MEM media containing RNPs at the concentration of 400ng/µl Cas9D10A and 400ng/µl of each sgRNA. The electroporation parameters were as follows: six poring pulses of 15V, with 10% decay, for 2ms with a 50ms interval, immediately followed by 5 transfer pulses of 3V, 40% decay, for 50ms with a 50ms interval, alternating the polarity. We conducted two electroporation sessions, the first at 14 hours post fertilization (hpf) and the second at 20 hpf. After the second electroporation, PZs were placed in culture media and incubated as indicated above.

We recorded the number of embryos that cleaved at ∼45hpf and blastocysts at ∼168hpf and ∼190 hpf. For statistical analysis we considered culture drops as biological replicates. We analyzed count data (success of blastocyst development or developmental arrest) using a general linear model with a binomial family, which results in logistic regression analysis (Cox 1958), using the “glmer” function from the R package “lme4” (Bates et al. 2015). We used the number of blastocysts and the number of putative zygotes that failed to develop into blastocysts as the dependent variable. Group (Cas9 + targeting gRNAs or Cas9 + scramble gRNAs) was a fixed effect and replicate was a random variable. The Wald statistical test (Wald 1945) was conducted with the function “Anova” from the R package “car” (Fox and Weisberg 2018). Finally, we carried out a pairwise comparison using the odds ratio and two-proportion z-test employing the “emmeans” function of the R package “emmeans”. The null hypothesis assumed that the odds ratio of the proportion (*p*) of two groups was not different from 1 (*H*_0_: *p*_1_/*p*_2_ = 1). We inferred significance when adjusted P value < 0.05.

To access the edits produced, at ∼190 hpf, we also collected all embryos that arrested their development at the morula stage. We removed their zona pellucida by a treatment with EmbryoMax Acidic Tyrode’s Solution (Millipore Sigma, Danvers, MA) and exposed their DNA by adding four µL of QuickExtract DNA Extraction Solution (Biosearch Laboratories, USA), and incubating the solution at 65°C for 15 min followed by 2 min at 98°C and hold at 4°C.

Then, we conducted PCR reactions using the oligonucleotides described on Supplementary table S15. The oligonucleotides were designed using NCBI’s Primer-BLAST (Ye et al. 2012) and certified their specificity using the University of California Santa Cruz’s BLAST-Like Alignment Tool available in the Genome Browser (Kent 2002; Kent et al. 2002). The PCR reaction mix consisted of 1.25 PrimeSTAR GXL DNA Polymerase (Takara Bio USA, San Jose, CA), 1× Buffer, 200 μM dNTPs, and forward and reverse oligonucleotides (IDT, Coralville, IA, USA) at 0.2 μM each, in a final volume of 50 μL. The cycling conditions for this reaction were: 98°C for 1 min, followed by 30 cycles of 98°C for 15 seconds, 62°C for 15 seconds, and 68°C for 4 minutes, followed by a final extension of 4 minutes at 72°C. We confirmed the amplification by assaying 5 µL of each amplicon by electrophoresis on a 1.5% Agarose gel before staining with Diamond Nucleic Acid Dye and imaging. Finally, we prepared libraries for amplicon sequencing with the Native Barcoding Kit 24 V14 (Oxford Nanopore Technologies, Lexington, MA, USA), following the manufacturer’s instructions. We sequenced the libraries in a MinION flowcell (R10.4.1) using a MinION Mk1C (Oxford Nanopore Technologies, Lexington, MA, USA) following the manufacturer’s recommendations.

We carried out super accuracy base calling with Guppy v6.4.2 (Wick et al. 2019), followed by quality filtering using Fitlong (https://github.com/rrwick/Filtlong) to remove short sequences (<500nt). Then, filtered reads were mapped to the cattle reference genome from Ensembl (ARS-UCD1.2) using minimap2 v2.27 (Li 2018, 2021) and sequences with <500nt aligned with the reference genome were removed by SAMtools (Li et al. 2009), as well as secondary alignments.

## DATA ACCESS

All raw and processed sequencing data generated in this study have been submitted to the NCBI Gene Expression Omnibus (GEO; https://www.ncbi.nlm.nih.gov/geo/) under accession identifiers GSE99678, GSE199210, GSE225693.

## ACKNOWLEDGEMENTS

The authors acknowledge Select Sires for the semen straw donation used in this project.

